# Single-cell RNA-sequencing of bronchoscopy specimens: development of a rapid, minimal-handling protocol

**DOI:** 10.1101/2023.03.02.530288

**Authors:** Firoozeh V Gerayeli, Stephen Milne, Chen Xi Yang, Xuan Li, Elizabeth Guinto, Julia Shun-Wei Yang, Chung Yan Cheung, Tara R Stach, Tawimas Shaipanich, Janice M Leung, Don D Sin

## Abstract

Single-cell RNA-sequencing (scRNA-seq) is an important tool for understanding disease pathophysiology, including airways diseases. Currently, the majority of scRNA-seq studies in airways diseases have used invasive methods (airway biopsy, surgical resection) which carry inherent risks and thus present a major limitation to scRNA-seq investigation of airway pathology. Bronchial brushing, where the airway mucosa is sampled using a cytological brush, is a viable, less invasive method of obtaining airway cells for scRNA-seq. Here we are describing the development of a rapid and minimal-handling protocol for preparing single cell suspensions from bronchial brush specimens for scRNA-seq. Our optimized protocol maximises cell recovery and cell quality, and may facilitate large-scale profiling of the airway transcriptome at single cell resolution.

**Lay abstract:** Single-cell RNA-sequencing (scRNA-seq) measures the gene expression of individual cells, and may be useful for understanding disease processes. scRNA-seq may be used to investigate lung diseases, but using invasive methods such as biopsy or surgery limits our ability to conduct large research studies. Bronchial brushing, where a soft brush is used to collect cells from inside the lungs, is a safer method but we need a better way to isolate individual cells from the brush specimens. We developed a method that is faster and involves less handling of the specimens compared to other published methods. Our method may therefore be useful for conducting large scRNA-seq studies in lung diseases.

## Introduction

The airway mucosa has a critical role in maintaining local and systemic homeostasis. Its protective functional properties include intercellular epithelial junctions, mucociliary clearance, soluble intraluminal immune factors, active immune surveillance, and initiation of innate and adaptive cellular immunity [1]. Profiling the transcriptome of the airway mucosa has revealed insights into normal airway physiology, as well as the pathophysiology of numerous diseases including asthma [2-4], chronic obstructive pulmonary disease (COPD) [5-8], and lung cancer [9,10]. However, bulk transcriptomics (either microarray or RNA-sequencing) is unlikely to have sufficient resolution to capture the complexity of the airway mucosal cellular landscape.

Single-cell RNA-sequencing (scRNA-seq) allows analysis of the transcriptome at a single cell resolution. The method uses microfluidic technology to capture single cells in a lipid droplet, where RNA transcripts are then reverse-transcribed – with an oligonucleotide label unique to each cell – within this local environment. Libraries are then amplified and sequenced using high throughput, next-generation RNA sequencing. The resulting, highly complex multi-dimensional data are computationally processed such that data points with similar transcriptomes are clustered together and annotated as cell populations. scRNA-seq of the airway mucosa has revealed enormous cellular heterogeneity [11] as well as novel cell types [12]. However, the majority of scRNA-seq data published to date have been derived from invasive airway biopsy, surgical resection, or post mortem specimens. Such specimens are difficult to obtain and highly invasive, and carry inherent safety risks. This presents a limitation to future large-scale scRNA-seq investigation of the airways, particularly in participants with comorbidities who have higher risk of complications.

Bronchial brush (BB) sampling of the airway mucosa is a less-invasive and a relatively safe procedure performed in routine clinical practice. A cytological brush is inserted, via a bronchoscope, into the airways and gently scraped against the airway wall to capture cells. In clinical practice, the specimens are assessed by cytology, microbiology, or nucleic acid tests. In research settings, BB specimens have been used for profiling the airway transcriptome [2,3,6-9] and microbiome [13]. More recently, a number of studies have used BB specimens for scRNA-seq analysis of the airways [14-19].

Despite many years of experience with research bronchoscopies and airway transcriptomics using BB specimens, our early attempts using previously published protocols for scRNA-seq were hampered by long processing times, low cell recovery, poor cell viability, and low RNA quality. We therefore developed a novel protocol for processing BB specimens for scRNA-seq, based on the following guiding principles: 1) minimize the handling and manipulation of cells, 2) create a single cell suspension with negligible cell clumping or debris; 3) maximize cell recovery and viability; and 4) minimize processing time. Here we describe the development of an optimized protocol for research. We also developed an optimized protocol for processing bronchoalveolar lavage (BAL) fluid based on the same principles. The resulting optimized protocols have facilitated a rapid expansion of our program with over 40 specimens processed in the past six months.

## Materials and Methods

### Human research ethics

The specimens used for this study were collected under protocols that were approved by the Human Research Ethics Board of the University of British Columbia/Providence Health Care (approval numbers H11-02713, H19-02222, and H21-02149). All study participants provided informed consent.

### Bronchial brush protocol

The details of our standardized research bronchoscopy protocol have been previously published [20]. After receipt of informed consent and under conscious sedation, a fibreoptic bronchoscope (Olympus Corporation, Tokyo, Japan) was passed through the mouth of participants and into their trachea. With the bronchoscope positioned in one of the subsegmental bronchus of the right (or left) upper lobe, a cytological brush was inserted through a bronchoscope channel into a 6^th^-8^th^ generation airway from where bronchial brush samples were collected. The cytologic brush was then withdrawn from the bronchoscope and using a pair of stainless steel scissors, the brush was cut into a microcentrifuge tube containing 1000 μL medium and kept on ice until further processing. We evaluated the following media for sample quality: Dulbecco’s phosphate-buffered saline (DPBS) (Gibco, Grand Island, NY, USA), RPMI 1640 (Corning, NY, USA) with and without fetal bovine serum (FBS) (Invitrogen, Waltham, MA, USA), and PneumaCult-Ex^™^culture medium (STEMCELL Technologies, Vancouver, BC, Canada).

For enzymatic digestion of intercellular junctions and extracellular matrix (ECM), approximately 1000 μL of Accutase^™^ (Gibco) were added directly to brush specimens in the medium, followed by gentle agitation. The digestion time was titrated from 3 to 10 minutes, and the enzyme activity was quenched by directly transferring the combined Accutase^™^/medium into a separate tube containing medium, either 2 mL (‘small volume protocol’) or 25 mL (‘large volume protocol’).

To wash the cells, the cell suspension was passed through a 70 μm cell strainer (VWR, Radnor, PA, USA) to remove large debris and cell clumps and centrifuged at 400g for 10 mins at 4 °C; after which the supernatant was discarded. Under the small volume protocol, the cell pellet was resuspended in 2 mL medium and centrifugation was repeated as above, whereas the large volume protocol included a single centrifugation step. The final cell pellet was resuspended in medium to a total volume of 100 μL, which was then filtered through either a 40 μm cell strainer (VWR) or a 40 μm Flowmi strainer (SP Bel-Art, Warminster, PA, USA).

### BAL protocol

BAL was performed by first wedging the bronchoscope into the target airway (preferentially the right middle lobe or the left lingular segment), instilling aliquots of warm saline into the distal lung and allowing it to dwell for 10 seconds, and then withdrawing the lavage fluid by suction. This process was repeated until 30 mL of BAL fluid was recovered, or a total instilled volume of 200 mL was reached, whichever came first. The recovered BAL fluid was then stored on ice until further processing.

The cellular content of BAL is almost exclusively single unattached cells; therefore, no enzymatic digestion was necessary to create a single cell suspension. The BAL fluid was first filtered through a 70 μm cell strainer to remove mucus and debris, then centrifuged at 400g for 10 minutes at 4 °C, and the supernatant was decanted and stored at -80 °C for future use. The cells were then washed by resuspending the pellet in 10 mL of DPBS + 0.04% (weight/volume) bovine serum albumin (BSA) (Invitrogen), centrifuging at 400g for 10 minutes at 4 °C, and discarding the supernatant. The final cell pellet was resuspended in 1,000 μL of DPBS + 0.04% BSA.

### Single-cell RNA sequencing

Visual assessments and cell counts of the final cell suspensions were performed using a haemocytometer with Trypan Blue stain (Gibco). When necessary, the cell suspensions were diluted to a concentration of 1,000 cells/μL in their respective media prior to single cell sequencing. For further examination, aliquots of the cell suspensions were applied to histological slides by cytospin and stained using a modified Wright-Giemsa stain.

### Cell capture and library preparation

Cell suspensions were transferred on ice to our sequencing facility (approximately 25 minutes away) and loaded onto the Chromium Controller or Chromium X using the Chromium Next GEM Single Cell 3’ Kit v3.1 and Chip G (10X Genomics). Samples were handled and libraries prepared according to the published Chromium Single Cell 3’ Reagent Kits User Guide [21]. The quality of the cDNA libraries and final DNA libraries were analyzed using the 2100 Bioanalyzer Instrument with the High Sensitivity DNA Kit (Agilent, Santa Clara, CA, USA).

### Sequencing

Final libraries were sequenced on the NextSeq2000 (Illumina, San Diego, CA, USA) as recommended by 10X Genomics (Read 1: 28bp, Read 2: 90bp, i7 index: 10bp, i5 index: 10bp), at a loading concentration of 650pM with a 2% PhiX Spike-in. All sequencing was initially performed at a shallow depth of 20,000 reads/cell, assessed, and then further sequencing performed until a depth of 60k reads/cell (BB specimens) or 40k reads/cell (BAL) was achieved. Fastq files were generated with Cell Ranger v6.01 (10X Genomics) using the default options, and reads were aligned to the human reference genome (hg19).

### Bioinformatics

A detailed description of the data processing is provided in the Online Supplementary Material (Figure S1). Analysis of the single cell RNA sequencing data can be challenging due to the presence of technical artifacts such as ambient RNA. Such contamination would inadvertently impact the downstream analysis of the samples. SoupX package version 1.5.2 [22] in R was used prior to performing individual quality control (QC) tasks to correct for ambient RNA, whose expression is projected from the empty droplet pool. Following ambient RNA correction, using Pegasus package version 1.5.0 [23,24] in Python, we performed additional QC steps, in which cells with high mitochondrial (MT) genes (MT genes >20%), and < 200 genes per cell were filtered out. This step was followed by log normalization, batch effect correction using the harmony algorithm, and doublet removal. For dimensionality reduction, we performed principal component analysis (PCA), prior to applying Uniform Manifold Approximation and Projection (UMAP) dimensionality reduction. Preceding to any further downstream analysis we annotate the dataset using Pegasus package automatic annotation, in conjunction with the markers noted in literatures.

## Results and Discussion

The central motivation for our work was to investigate the pathology of airway diseases using scRNA-seq. Bronchial brushing is an attractive method of sampling the lung because it is relatively non-invasive, safe, and is routinely performed in clinical practice. A number of investigators have reported scRNA-seq using BB specimens [14-19]. However, a review of these published protocols revealed key differences in the way the specimens were processed (Table 1). Additionally, the number of cells recovered per participant was highly variable and none of the published protocols were accompanied by detailed validation of their results. We therefore sought to develop our own protocol that maximized cell recovery and quality. The final protocols for BB and BAL specimens are included in the Online Supplementary Material (“BronchialBrush_Protocol” and “BAL_protocol”, respectively).

**Table 1:**
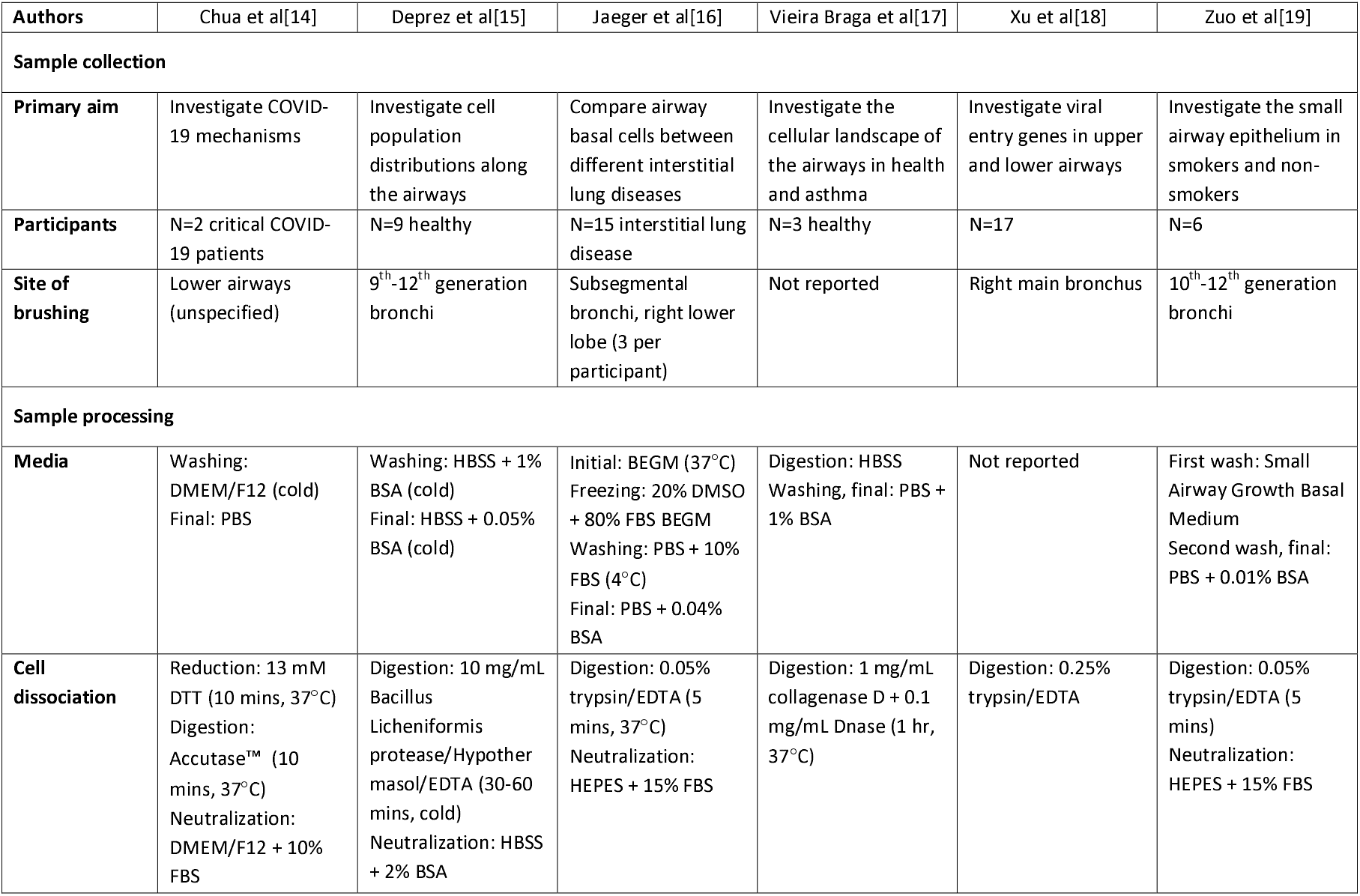
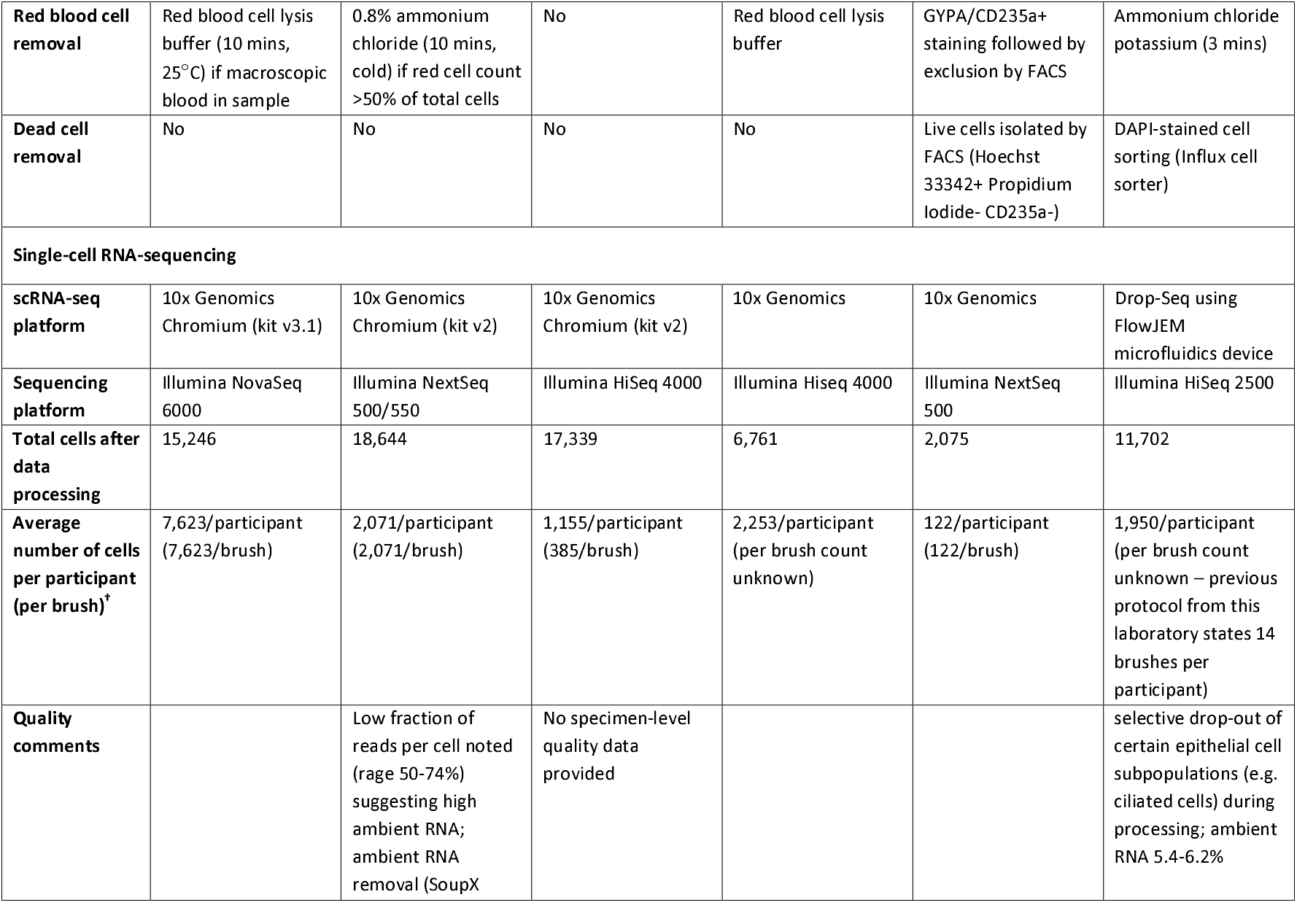
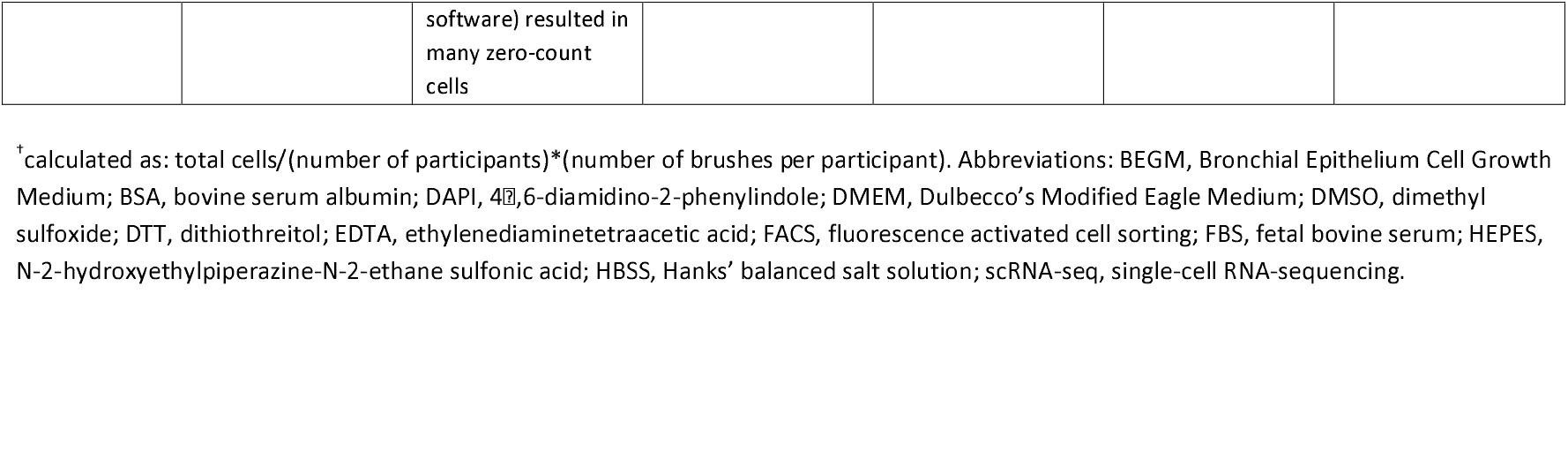
Summary of published protocols for bronchial brush processing for single-cell RNA-sequencing

The choice of medium is important for not only maintaining cell viability but also for supporting single cell suspension. The 10X Genomics standard protocol [25] recommends a calcium/magnesium free balanced salt solution spiked with protein, which minimises cell losses and aggregation. Indeed, the published BB protocols used either FBS or BSA (Table 1). However, we found that the use of these solutions for BB specimens was associated with poor cell viability and excessive clumping of cells, which led to low concentrations of viable cells at the end of processing. We found that the best medium was Pneumacult-Ex^™^, a serum- and bovine pituitary extract-free cell culture medium specifically formulated to support the growth of airway epithelial cells. We used this medium at all stages of processing BB specimens in the optimized protocol. To our knowledge, this is the first time this medium has been used for scRNA-seq experiments.

In order to generate a single cell suspension from BB specimens, it was necessary to incubate the brushes in a protease solution. Enzymatic digestion of whole tissue and biopsy specimens is routinely used to break down ECM and create single cell suspensions. Although BB specimens contain little ECM, the epithelial cells scraped from the airways were connected by intercellular junctions and therefore tended to remain in sheet-like clumps (Figure 1). The use of protease treatment of BB specimens was common to all previously published protocols, though most employed non-specific proteases such as trypsin (Table 1). We elected to use Accutase^™^, an enzyme mixture with proteolytic and collagenolytic properties, since it is associated with efficient cell dissociation without interrupting cell surface antigens or inducing cell cycle changes [26], and leads to improved viability when culturing fragile cell types [27]. Digestion for 10 minutes (as used by Chua et al) [14] resulted in widespread cell destruction and fragmentation, with occasional single cells which lacked the expected morphology of bronchial epithelial cells (Figure 2a). After single cell capture and cDNA library preparation, the bioanalyzer indicated high levels of short sequences suggestive of either high ambient RNA or fragmented RNA in the specimens (Figure 2b). The processed data confirmed high levels of ambient RNA, a low number of cells (far below the expected number based on the sample input), and only three identifiable cell clusters (Figure 2c). Notably, the proportion of epithelial cells was very low (Figure 2d), suggesting that this cell type is particularly sensitive to the effects of prolonged protease activity. In contrast, Accutase^™^ digestion for 3 minutes produced a single cell suspension with readily-identifiable bronchial epithelial cells including ciliated and goblet cells, as well as immune cells such as monocytes, and the resulting cDNA bioanalyzer output showed superior quality of RNA (Figure 3a). The processed scRNA-seq data demonstrated cell numbers closer to the expected count, with a diverse population of epithelial and immune cells consistent with the expected composition of the airway mucosa (Figure 3b and 3c). We therefore concluded that 3 minutes was the optimal duration of Accutase^™^ incubation.

**Figure 1:**
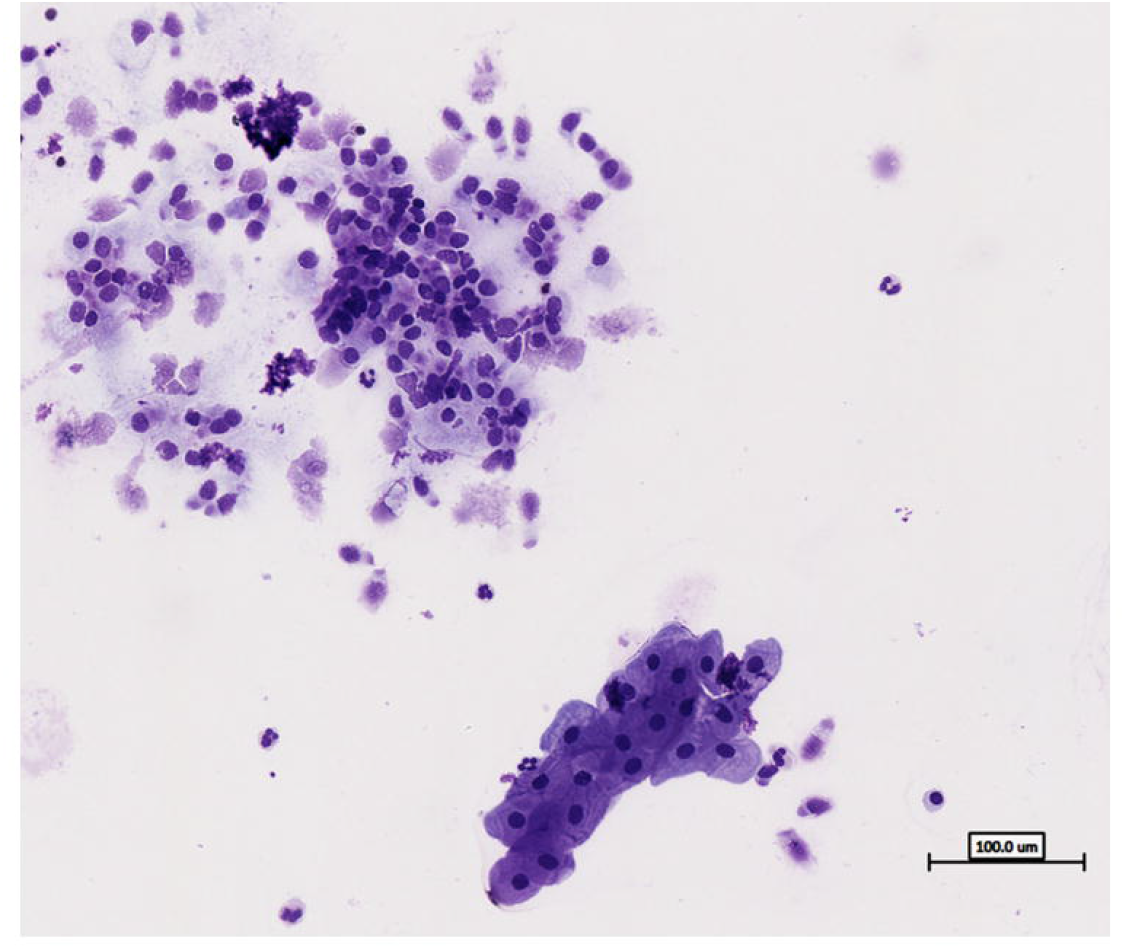
Bronchial brush specimens, untreated. Cells were retrieved by bronchial brushing from a single airway. Brush was immediately placed in Pneumacult-Ex^™^medium, gently vortexed, and centrifuged at 400g for 10 minutes at 4 degrees Celsius. The majority of cells were epithelial cells in large sheet-like clumps, with scattered immune cells and acellular debris. Modified Wright-Giemsa stain, 40X magnification.

**Figure 2:**
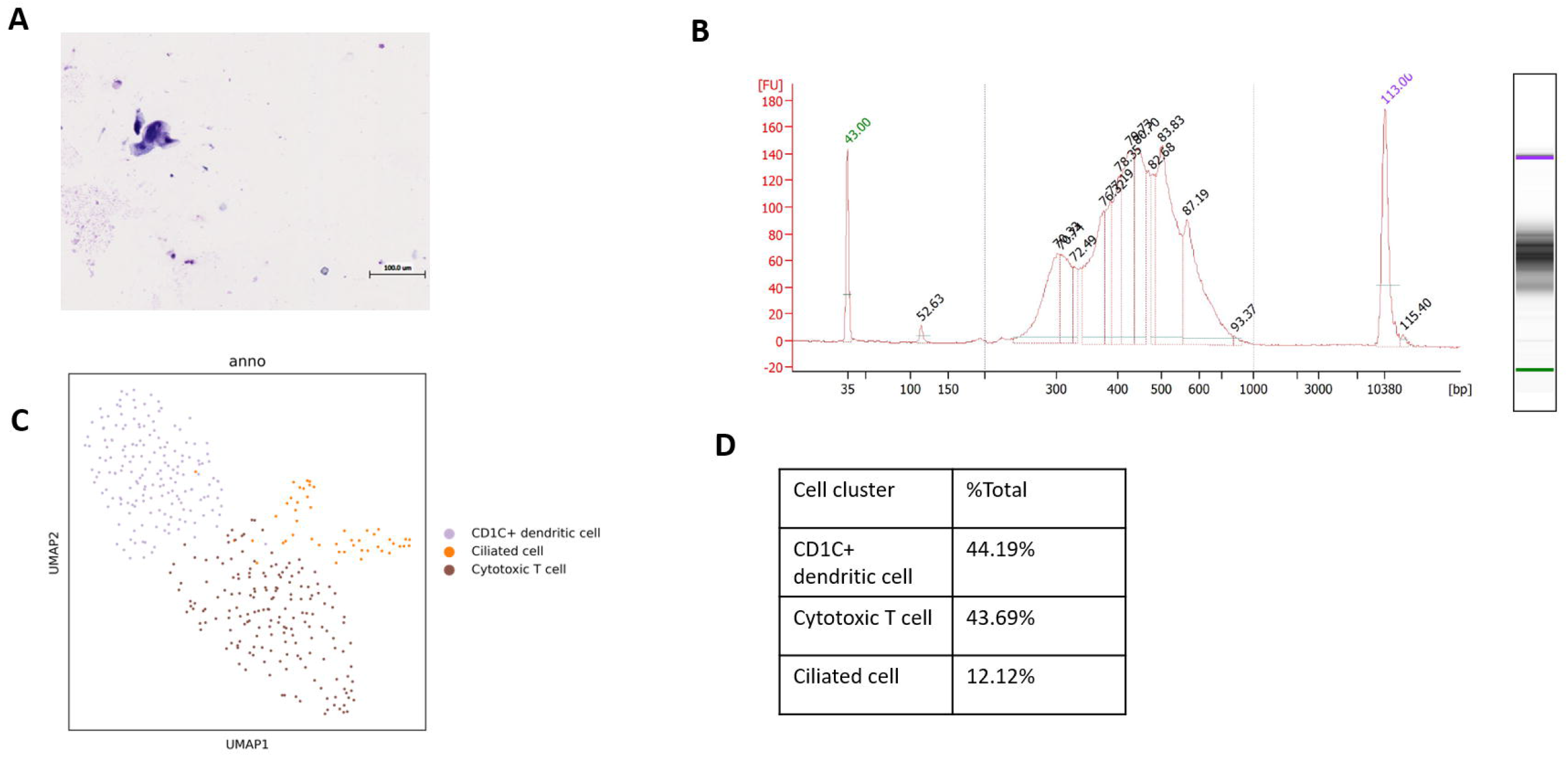
Bronchial brush specimens, 10 minutes Accutase^™^ digestion. 1 mL Accutase^™^ was added to the brush in medium for 10 minutes on a plate shaker. Digestion was stopped by adding an excess of medium. Sample was then centrifuged at 400g for 10 minutes at 4 degrees Celsius. (A) Modified Wright-Giemsa stain, 40X magnification, showing widespread cell destruction and fragmentation and occasional small cell that lack the known epithelial cells morphology. (B) Bioanalyzer output of the cDNA library following cell capture and RT-PCR, showing high number of short sequences suggestive of RNA fragmentation. (C) Uniform manifold approximation and projection (UMAP) of the scRNAseq data showing three cell clusters. (B) Cell proportions from scRNAseq, with a minority annotated as epithelial cells

**Figure 3:**
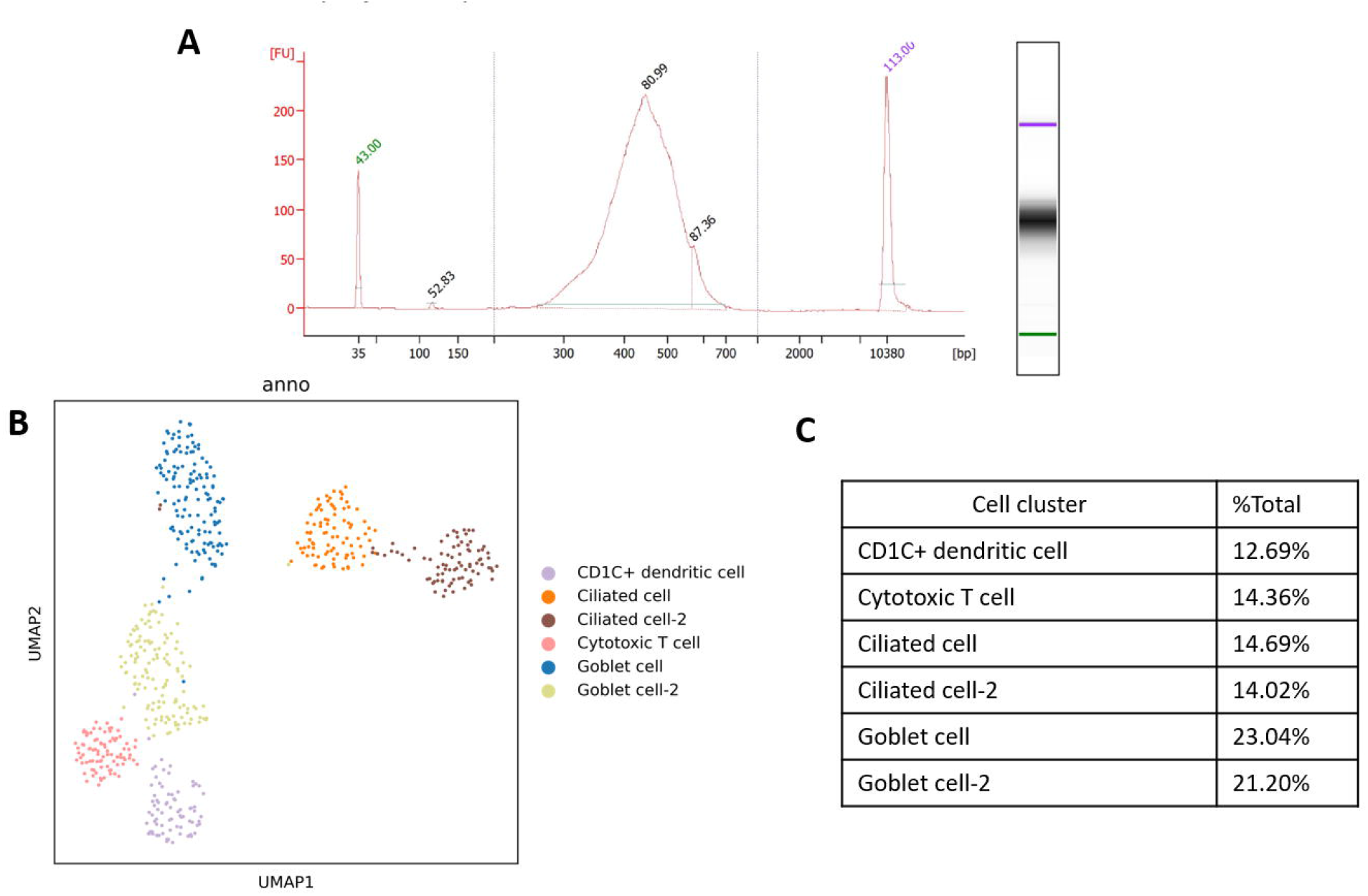
Bronchial brush specimens, 3 minutes Accutase^™^ digestion. 1 mL Accutase^™^ was added to the brush in medium for 3 minutes on a plate shaker. Digestion was stopped by adding an excess of medium. Sample was then centrifuged at 400g for 10 minutes at 4 degreesCelsius. (A) Bioanalyzer output of the cDNA library following cell capture and RT-PCR. (B) Uniform manifold approximation and projection (UMAP) of the scRNAseq data showing multiple cell clusters of both epithelial and immune cell lineages. (C) Cell proportions from scRNAseq.

Washing with medium followed by centrifugation not only removes ambient (i.e. extracellular) RNA but also eliminates residual proteases from the cells. The previously published protocols for BB specimens, as well as the 10X Genomics standard protocol [25], employed a minimum of two wash-centrifuge cycles (Table 1). However, we found that multiple wash cycles led to excessive cell loss. We therefore employed a single, large volume wash of 25 mL Pneumacult-Ex^™^ for BB specimens, which greatly improved the cell yield whilst minimising ambient RNA. We also removed debris and cell clumps using mechanical filtration. The presence of cell clumps not only leads to multiplets in the sequencing output (i.e. more than one cell being captured in an oil droplet) but also interferes with the microfluidics technology and generation of the gel emulsion [25]. We found that a 70 μm cell strainer was adequate to remove large clumps and debris such as mucus as an initial step. However, the cell strainer required adequate rinsing with clean medium to minimise the loss of single cells captured by the strainer. Since cell retrieval from BB specimens was typically low, this use of extra medium further diluted our cell concentrations. In order to minimize the volume and therefore increase cell concentration, we used a 40 μm Flowmi^™^ cell strainer as the final filtration step. This filter attaches directly to the pipette tip and can be used to filter volumes as low as 50 μL with minimal volume loss. In our experience, we were able to filter volumes of 100-120 μL with a typical volume loss of 20-40 μL.

To optimize the BB specimen protocol, there were several other considerations. The 10x Genomics standard protocol [25] and some of the published protocols [18,19] employed a dead cell removal step (e.g. by flow cytometry) in order to maximise the viable cell concentration. However, we elected to forgo this step for three reasons: first, our cell recovery was typically low and dead cell removal would further reduce the final cell concentration; second, we considered our cells fragile and needed to avoid further mechanical stress; and third, the quality of our cells was highly dependent on the time since collection, meaning delays related to this step would further compromise the overall quality of the cells. We also did not employ red blood cell lysis techniques since our bronchial epithelial cells were sensitive to the typical lysis reagents (e.g. ammonium chloride); instead, we avoided using macroscopically-contaminated specimens and removed any red blood cell ‘contaminants’ in silico using bioinformatics tools. We kept specimens and reagents at 4 °C rather than at room temperature (as recommended by the 10X Genomics standard protocol) [25], because in our experience this minimised cell clumping. Finally, we used low retention materials (conical and microcentrifuge tubes) wherever possible to maximise cell recovery. With each of these measures, our BB specimen protocol produced high-quality, single cell suspensions with increased and reliable cell counts (Table 2). The processed sequencing data showed that all the expected cell types in the airway epithelium were present (Figure 4).

**Table 2:** Bronchial brush protocol optimization and outcomes

| **Participant characteristics** | 75 y/o female with lung mass | 42 y/o female with asthma | 60 yr old male with COPD | 59 yr old female with COPD | 62 yr old male with COPD |
| --- | --- | --- | --- | --- | --- |
| **Sample preparation** |  |  |  |  |  |
| **Medium** | DPBS | DPBS +10% FBS | RPMI1640 | Pneumacult-Ex™ | Pneumacult-Ex™ |
| **Temperature** | Room temperature | Room temperature | Room temperature | Specimens and reagents kept on ice; centrifuged at 4°C | Specimens and reagents kept on ice; centrifuged at 4°C |
| **Cell dissociation** | Accutase™ | Accutase™ | Accutase™ + plate shaker | Accutase™ + gentle agitation | Accutase™ + gentle agitation |
| **Protease incubation time** | 10 minutes | 10 minutes | 10 minutes | 3 minutes | 3 minutes |
| **Wash-centrifuge steps** | 1 x 5 mL wash | 3 x 5 mL wash | 1 x 25 mL wash | 1 x 25 mL wash | 1 x 25 mL wash |
| **Filtration** | 40 µm bucket strainer | 40 µm bucket strainer | 40 µm bucket strainer | 70 µm and 40 µm bucket strainers | 70 µm bucket and 40 µm Flowmi™ strainers |
| **Final volume** | 150 µL | 150 µL | 100 µL | 100 µL | 100 ul |
| **Final cell concentration (hemocytometer)** | Overall non-viable cells – not counted | Overall non-viable cells – not counted | 500 cells/µL | 333 cells/µL | 505 cells/µL |
| **Morphology comments** | Majority dead/fragmented cells, residual debris | Majority dead/fragmented cells | Abnormal cell morphology (small, round); some cell clumping | Ciliated, other epithelial, and immune cell morphologies observed | Ciliated, other epithelial, and immune cell morphologies observed |
| **Single-cell RNA-sequencing** |  |  |  |  |  |
| **scRNA-seq platform** | Not submitted | Not submitted | 10x Genomics Chromium (kit v3.1) | 10x Genomics Chromium (kit v3.1) | 10x Genomics Chromium (kit v3.1) |
| **RNA sequencing** | Not submitted | Not submitted | Illumina NextSeq | Illumina NextSeq | Illumina NextSeq |
| **Number of cells called** | Not submitted | Not submitted | 572 | 1,017 | 1,102 |
| **Data quality comments** | Not submitted | Not submitted | RNA fragmentation/high ambient RNA; low fraction of reads in cells (39%) | Low fraction of reads in cells (66%) | Low fraction of reads in cells (66.4%) |

Abbreviations: COPD, chronic obstructive pulmonary disease; DPBS, Dulbecco's phosphate buffered saline

**Figure 4:**
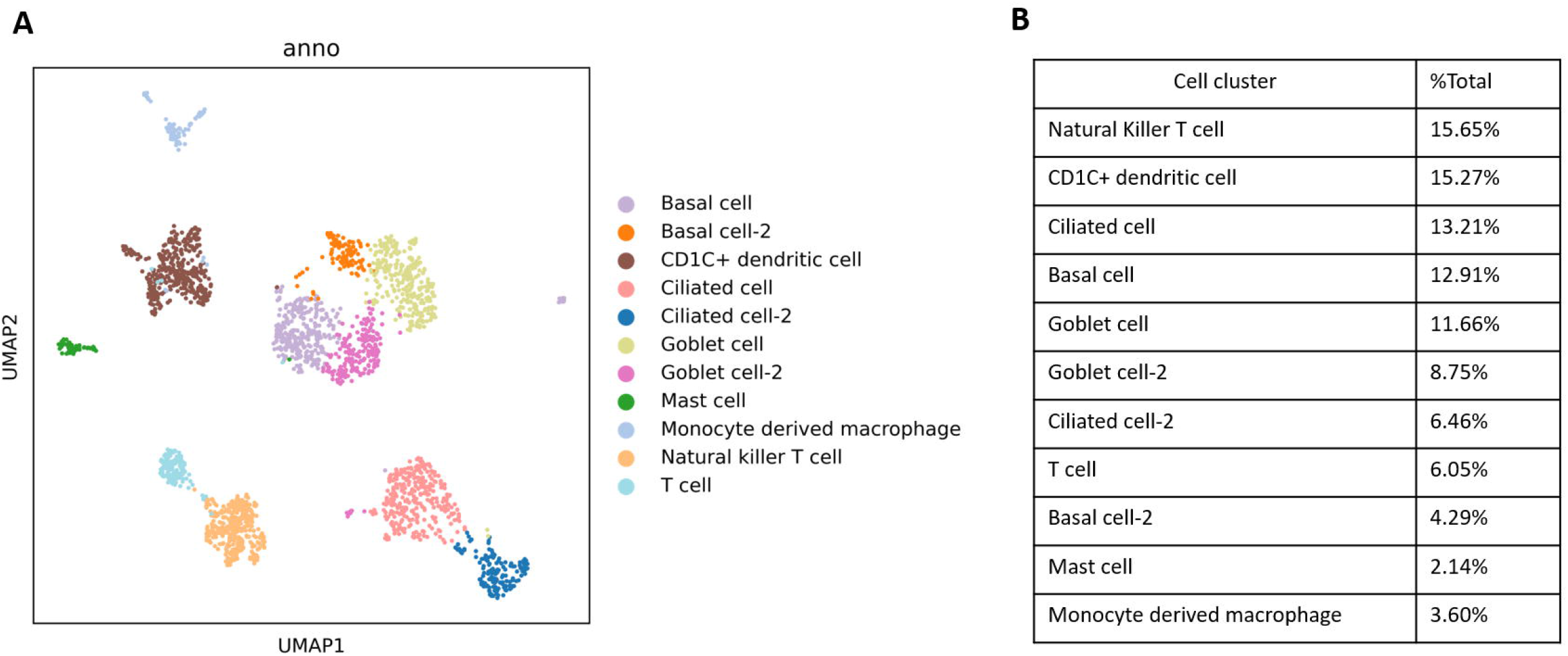
(A) Uniform manifold approximation and projection (UMAP) of the scRNAseq data of 2 subjects showing various cell clusters. (B) Cell proportions from scRNAseq, consist of annotated epithelial and immune cells

For the sake of completeness, we have included our optimized BAL processing protocol (Online Supplementary Material, “BAL_protocol”). Our protocol was derived from the 10X Genomics standard protocol, but we made three important modifications based on our experience with BB specimens. First, we used a large pore (70 μm) cell strainer as an initial filtration step to remove large debris (predominantly mucus). Depending on the sample, if the strainers became clogged with mucus; we advocate changing to a fresh filter rather than forcing the specimen through, to avoid mucus contamination in the cell suspension. Second, we used large volume washes (10 mL) to help minimise debris and ambient RNA. Third, we used a higher centrifuge speed of 400g for 10 minutes; in our experience with BAL fluid, this speed leads to optimal cell recovery and viability whilst minimizing debris. Our scRNA-seq output for BAL shows excellent quality cDNA libraries, minimal ambient RNA, and cell numbers that are consistent with the known specimen concentrations (data not shown). A representative UMAP of a BAL from a single subject submitted for scRNA-seq is shown in Figure S2.

## Conclusion

We developed a protocol for preparing BB and BAL specimens for scRNA-seq that maximises cell viability and recovery by minimizing specimen handling and processing time. The use of this optimized protocol has allowed us to build a scRNA-seq dataset from over 50,000 cells representing airways of 26 participants (BB) and over 100,000 cells from the BAL of 24 participants. This large dataset will be used to explore the pathology of various airways diseases, and how changes in the airways at single-cell resolution may relate to clinically important outcomes.

## Summary points

- Single-cell RNA-sequencing (scRNA-seq) allows analysis of the transcriptome at a single cell resolution.
- scRNA-seq reveals heterogeneity in the airway cellular landscape.
- Currently, the majority of data published on scRNA-seq have been derived from surgical biopsies, surgical resection of tissues including explants, or post mortem specimens.
- Bronchial brush sampling of the airway mucosa may be a safer, less-invasive alternative method to investigate airways.
- We have developed a protocol for creating single cell suspensions from bronchial brush samples with negligible cell clumping and debris, while minimizing the processing time.
- We also developed an optimized protocol for processing bronchoalveolar lavage specimens for scRNA-seq, which minimizes processing time and maximizes cell recovery.
- Our protocols will facilitate and standardize large-scale scRNA-seq experiments, enabling new insights on the pathophysiology of airway diseases.

## Supporting information

Bronchoalveolar Lavage Processing SOP

Bronchial Brush Processing SOP

## ACKNOWLEDGEMENTS

FVG is supported by Mitacs. EG is supported by Mitacs. SM is supported by Mitacs and Michael Smith Health Research BC. JML is a Tier 2 Canada Research Chair. DDS holds a Tier 1 Canada Research Chair in COPD and the de Lazzari Family Chair at HLI.

## Figure legends

S1: A detailed description of bronchial brush and BAL data processing pipeline

S2: Uniform manifold approximation and projection (UMAP) of the scRNAseq data of a 62 year old male with chronic obstructive pulmonary disease showing multiple cell clusters of immune cells and few epithelial cells

