## Supplementary figures and images for "Single-cell RNA-sequencing of bronchoscopy specimens: development of a rapid, minimal-handling protocol"

### Supplemental Figure 1

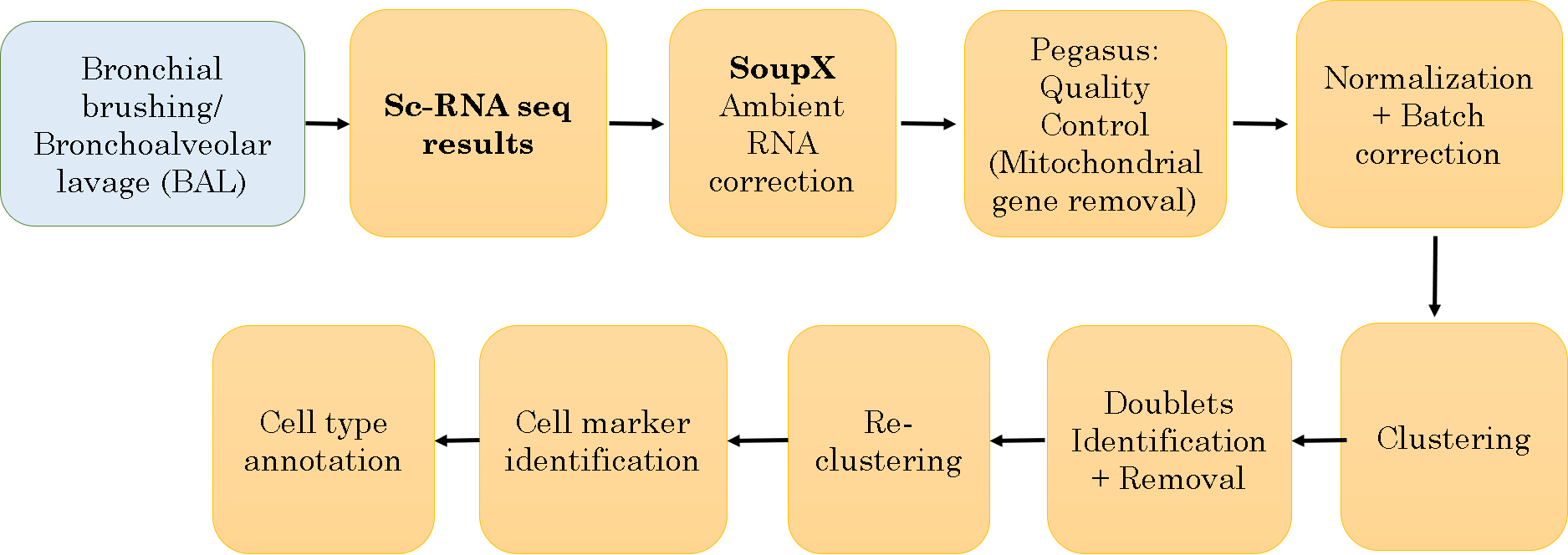

### Supplemental Figure 2

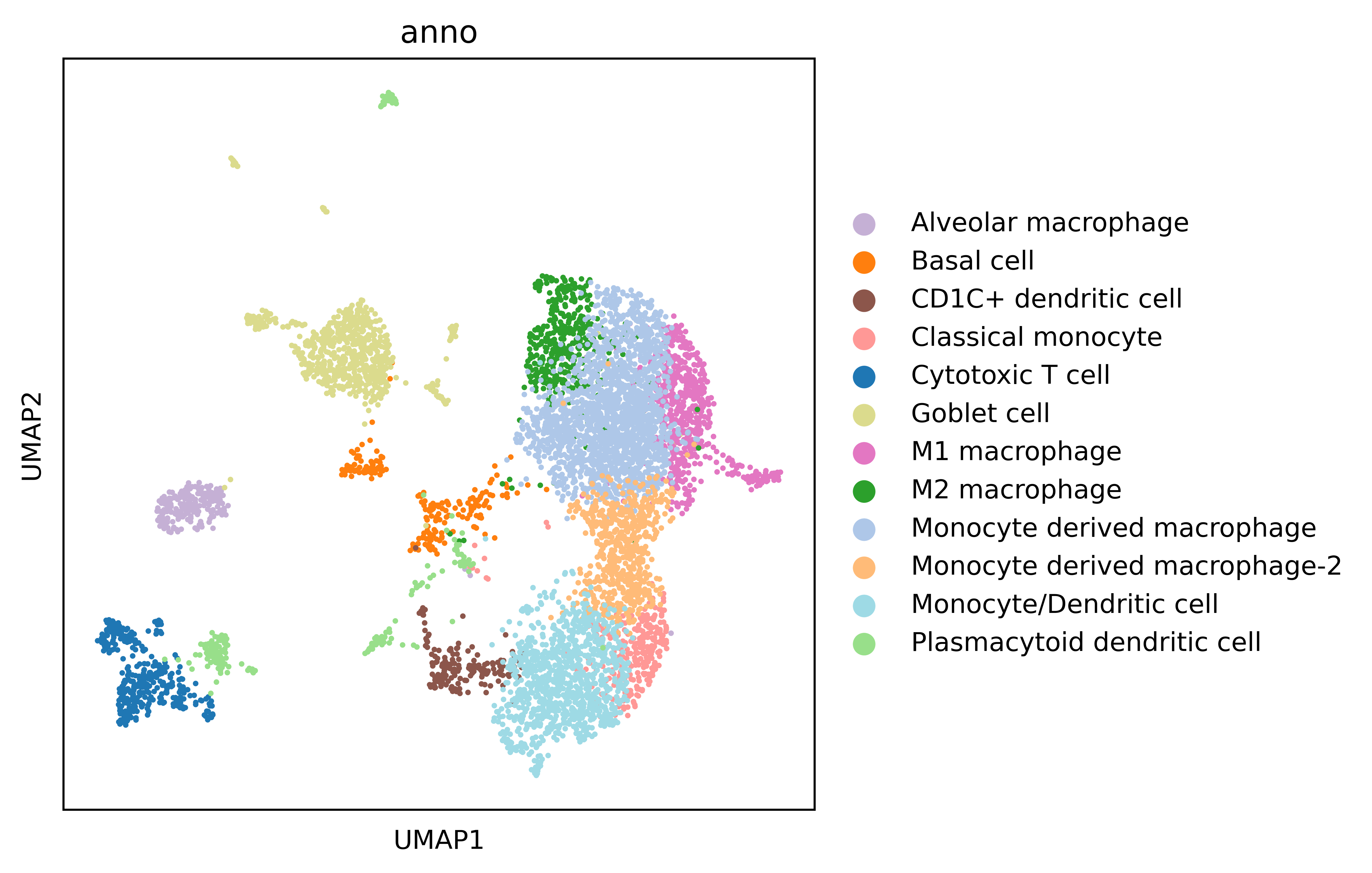
