## Supplementary material for "Single-cell RNA-sequencing of bronchoscopy specimens: development of a rapid, minimal-handling protocol": Bronchoalveolar Lavage Processing SOP

**PROTOCOL FOR:**

**SINGLE-CELL SUSPENSIONS FROM BRONCHOALVEOLAR LAVAGE (BAL) SPECIMENS**

### LEGEND

🢡***ATTENTION***	* ***HINT***

### REAGENTS AND MATERIALS

LoBind 50 mL DNase/RNase free conical tube (Eppendorf, Hamburg, Germany)
CELLSTAR serological 10ml DNase/RNase free pipette (Greiner, Kremsmünster, Austria)

70 μm bucket cell strainer (VWR, Radnor, Pennsylvania)

Dulbecco's phosphate-buffered saline (DPBS) (Gibco, Waltham, Massachusetts, USA)

Ultra Pure’ RNase free BSA (50 mg/mL) (ThermoFisher, Waltham, Massachusetts, USA)

LoBind 1.5 mL DNase/RNase free microcentrifuge tubes (Eppendorf ,Hamburg, Germany)

Wide Bore Hydrophobic DNase/RNase free Filter Barrier Pipet Tips, Axygen Scientific (1000 uL) (Union City, California)

Trypan Blue solution(Gibco, Waltham, Massachusetts, USA)

### PROCEDURE

1) 🢡All processing steps should be done in biosafety cabinet
2) Strain the BAL through the 70 µm bucket cell strainer into a 50 mL LoBind conical tube

*If strainer becomes clogged, tilt the strainer to use the side mesh. Use a new strainer if necessary.

🢡If the BAL is contaminated with blood it is advisable to not submit the sample for single cell RNA sequencing

3) Centrifuge @ 400g, 4°C, 10min
    🢡If the pellet contains blood it is advisable to not submit the sample for single cell RNA sequencing
4) Decant the supernatant

🢡To minimize ambient RNA contamination, remove as much of the supernatant as possible. This may require pipetting any remaining media from the sides of the conical tube.

5) Gently resuspend the cell pellet in 10 mL of DPBS + 0.4% BSA using 100p wide bore pipette
6) Centrifuge @ 400g, 4°C, 10min
7) Decant the supernatant
8) Dislodge the pellet by gently flickering the bottom of the conical tube and then using wide bore 1000p pipette. Gently resuspend in DBPS + 0.4% BSA up to a final volume of 500 µL (if estimated cell number is <500k) or 1000 µL (if estimated cell number is >500k)

*An estimate of the total cell number can be made from the cell pellet – a clearly-visible, white, solid looking pellet is likely to have >500k cells whereas a pellet that is thin and barely visible is likely to have <500k cells. If in doubt, resuspend in the smaller volume (500 µL) and then dilute if the cell concentration is too high to count confidently.

9) Resuspend in Trypan Blue (final 0.2%) and perform cell count with a hemocytometer under a microscope

🢡Count only viable (i.e. unstained) cells, and do not count red blood cells

10) Dilute the specimen to the optimum cell concentration (see recommendations for the single cell sequencing platform to be used). Transfer the cell suspension to a LoBind microcentrifuge tube ready for single cell capture.

### RECIPES

24.8ml DPBS + 200ul BSA

### EQUIPMENT

Eppendorf c 5810 R

Hemocytometer

Microscope
