## Supplementary material for "Single-cell RNA-sequencing of bronchoscopy specimens: development of a rapid, minimal-handling protocol": Bronchial Brush Processing SOP

### PROTOCOL FOR:

**SINGLE-CELL SUSPENSIONS FROM BRONCHIAL BRUSH SPECIMENS**

#### LEGEND

🢡***ATTENTION***	* ***HINT***

### REAGENTS AND MATERIALS

### Accutase (Gibco Waltham, Massachusetts, USA,)

PneumaCult-Ex culture medium (STEMCELL Technologies, Vancouver, BC, Canada)
Trypan Blue stain (Gibco, Waltham, Massachusetts, USA)

Flowmi 40 μm strainer (SP Bel-Art, Warminster, PA, USA)
40 μm bucket cell strainer (VWR, Radnor, Pennsylvania)
Wide Bore Hydrophobic DNase/RNase free Filter Barrier Pipet Tips, Axygen Scientific (1000 uL) (Union City, California)
Wide Bore Hydrophobic DNase/RNase free Filter Barrier Pipet Tips, Axygen Scientific (200 uL)
CELLSTAR serological 10ml DNase/RNase free pipette (Greiner, Kremsmünster, Austria)

Eppendorf Lobind 50 mL DNase/RNase free conical tubes (Hamburg, Germany)

Eppendorf Lobind 5 mL DNase/RNase free microcentrifuge tubes (Hamburg, Germany)

PROCEDURE

1) 🢡All processing steps should be done in biosafety cabinet

### 2) After collecting airway cells, place the cytological brush in a 5 mL microcentrifuge tube containing 1000 mL of cold PneumaCult-Ex media

🢩	Batch of used PneumaCult-Ex media should be prepped fresh every time according to the manufacturer instructions.

🢩	Airway cells are fragile, and the brush specimens should be processed without delay
🢩 If the brush is contaminated with blood it is advisable to not submit the sample for single cell RNA sequencing

### 3) Pulse vortex the brush 3x for 2 seconds

### 4) Gently invert the microcentrifuge tube containing the brush for 10X

### 5) Add 1000 µL of room-temperature Accutase to the microcentrifuge tube and mix by gently pipetting 2-3 time. Incubate on the plate tilter for 3 min (tilt: 15, speed 20)

### 🢩 Advisable to use 1000p wide bore DNAse/RNAse free pipette tips for this step.

* If there is excessive mucus in the specimen, gently pipette up to 10x to help break up the mucus prior to incubation

### 6) Quench the digestion by using 1000ul wide bore pipette tip for transferring the PneumaCult-Ex/Accutase mixture directly into a conical tube containing 25 mL fresh PneumaCult-Ex media. Wash the microcentrifuge tube/brush with at least 2 mL of PneumaCult-Ex media, and transfer the contents into the conical tube

### 7) Pour the media/cells from the conical tube through a 40 µm bucket cell strainer into a new conical tube. Wash the empty tube with 5 mL of fresh media and pass the contents again through the 40um bucket cell strainer.

### 8) Centrifuge at 400g for 10 mins at 4 ºC 🢡If the pellet is contaminated with blood it is advisable not to submit the sample for single cell RNA sequencing.

### 9) Decant and discard the supernatant

🢡To minimize ambient RNA contamination, remove as much of the supernatant as possible. This may require pipetting any remaining media from the sides of the conical tube.

### 10) Gently resuspend the pellet using a 200 µL wide bore pipette tip, without adding any extra media

🢡Pipetting must be slow and gentle in order to not damage the cells or introduce bubbles

### 11) Pass the resuspended cells through a Flowmi cell strainer attached to the pipette tip

* Take note of the final volume of the cell suspension – this will be necessary to determine the number of cells submitted for single-cell RNA sequencing

### 12) Perform a cell count using Trypan Blue (1:1 dilution) and hemocytometer

### RECIPES

Per manufacturer’s instruction:
24.5ml Pneumacult-EX media base + 500ul Pneumacult-EX media supplement+ 25ul hydrocortisone stock solution

### EQUIPMENT

Eppendorf Centrifuge 5810 R

VWR Rocking platform shaker

Hemocytometer

Microscope
